# Mapping language and non-language cognitive deficits in post-stroke anomic aphasia

**DOI:** 10.1101/2021.02.15.431293

**Authors:** Haya Akkad, Thomas M.H. Hope, Charlotte Howland, Sasha Ondobaka, Katerina Pappa, Davide Nardo, John Duncan, Alexander P. Leff, Jenny Crinion

## Abstract

While language impairment is the defining symptom of aphasia, the co-occurrence of non-language cognitive deficits and their importance in predicting rehabilitation and recovery outcomes is well documented. Despite this, people with aphasia (PWA) are rarely tested on assessments of higher order cognitive functions, making it difficult for studies to associate these functions with a consistent lesion correlate. Contrary to classic models of speech and language, cumulative evidence shows that Broca’s area and surrounding regions in the left inferior frontal cortex (LIFC) are involved in, but not specific to, speech production – suggesting that these regions may be involved in higher-level cognitive functions that support language production. A better understanding of language processing in the context of other domain general cognitive functions is essential for improving aphasia treatments.

This study aimed to explore the brain-behaviour relationships between tests of individual cognitive skill and language abilities in people with post-stroke aphasia, with a focus on language production deficits and their associated lesion correlates. We predicted our analysis would reveal a latent (non-language specific) cognitive component, that would be driven by damage to LIFC.

We analysed the behavioural and neural correlates of an extensive battery of language and non-language cognitive tests in a sample of thirty-six adults with long-term speech production deficits from post-stroke aphasia. All participants were anomic, with relatively intact speech comprehension and no apraxia of speech. The behavioural variables were analysed using Principal Component Analysis and their neural correlates were estimated using Voxel-Based Correlational Morphology. A significant number of anomic adults showed impaired performance on tests of non-language specific cognitive function. The variance underlying behavioural performance was best captured by four orthogonal components, two higher-order cognitive components (executive functions and verbal working memory) and two linguistic processing components (phonology and semantics). Brain-behaviour relationships revealed separable neural correlates for each component in line with previous studies and an executive functions correlate in the left inferior frontal cortex (LIFC).

Our findings suggest that in adults with chronic post-stroke language production deficits (anomia), higher-level cognitive functions explain more of the variance in language function than classical models of the condition imply. Additionally, lesions to the LIFC, including Broca’s area, were associated with executive (dys)function, independent of language abilities, suggesting that lesions to this area are associated with non-language specific higher-level cognitive functions that support speech production. These findings support contemporary models of speech production that place language processing within the context of domain-general perception, action and conceptual knowledge.

## 1. Introduction

While language impairment is the defining consequence of post-stroke aphasia, the presence of co-occurring impairments in other cognitive domains has been well documented (Fucetola et al., 2009; Helm-Estabrooks, 2002; Murray, 2012; El Hachioui et al., 2014; Marinelli et al., 2017; Ramsey et al., 2017; Schumacher et al., 2019) Despite this, People With Aphasia (PWA) rarely receive extensive cognitive assessment, meaning data on individual cognitive skills in this patient population is scarce. Evidence suggests that executive functions may be impaired in post-stroke aphasia, but the relationship between language and executive functions is difficult to tease apart (see Fedorenko, 2014) and studies have not been able to converge on the underlying lesions correlates of executive functions in PWA (Mirman and Thye, 2018).

Deficits in non-language specific cognitive domains have consistently been shown to be predictive of certain aspects of language function recovery in post-stroke aphasia. Marinelli and colleagues (2017) examined language and cognitive function in 189 PWA and found more severe language deficits to be associated with more severe cognitive impairments. Other studies have investigated executive functions in PWA and consistently found impaired inhibition, working memory or cognitive flexibility (Frankel et al., 2007; Fridriksson et al., 2006; Jefferies, Patterson and Ralph, 2008; Lee and Pyun, 2014; Murray, 2012; Vallila-Rohter and Kiran, 2013). A better understanding of language processing in the context of other domain-general cognitive functions is important for clinical management and rehabilitation. In fact, a series of aphasia therapy studies emphasise that cognitive abilities, particularly executive functions and verbal short-term memory, play an important role in driving recovery outcomes (Fillingham, Sage and Lambon Ralph, 2005a, 2005b, 2006; Conroy, Sage and Lambon Ralph, 2009; Mirman et al., 2015; Lambon Ralph et al., 2010; Yeung and Law, 2010; Snell, Sage and Lambon Ralph, 2010, Sage, Snell and Lambon Ralph, 2011; Dignam et al., 2017; Lacey et al., 2017; Schumacher et al., 2020).

While studies have highlighted the impact of cognition on aphasia rehabilitation and recovery, few have explored the contribution of individual cognitive skills and the relationship to underlying lesion pattern. The neural basis of aphasia is commonly explored by linking behavioural assessment with brain lesion data. This has resulted in some distinct brain-behaviour relationships for various language domains, however studies have not been able to converge on a consistent lesion correlate of higher-level executive functions (Mirman and Thye, 2018), either because non-language assessments were not included (Kummerer et al., 2013; Mirman et al., 2015) or were only included in a limited scope (Butler et al., 2014; Halai et al., 2017; Tochadse et al., 2018; though see Lacey et al., 2017). More recently, the neural correlates of non-language cognitive domains in aphasia have been explored by Schumacher et al., (2019, 2020) and Alyahya et al., (2020), whose findings are discussed in more detail below.

When assessing cognitive abilities, it is important to consider that cognition is a multidimensional construct broadly comprising five general domains, including language, attention, memory, executive functions and visuo-spatial skills (Helm-Estabrooks, 2002), with each domain containing distinct components. Using composite or general scores risks reducing the sensitivity of the cognitive measure. Schumacher and colleagues (2019) recently demonstrated the importance of this by using a detailed non-verbal neuropsychological assessment to show that brain regions involved in particular components of the attention and executive functions domains contribute to the abilities of adults with a wide range of aphasia types. Lacey and colleagues (2017) showed that executive functioning explains considerable variance in language abilities of PWA. Schumacher et al., (2020) recently showed that variance in functional communication abilities in PWA can be almost entirely explained by patients’ verbal short-term memory. Another study used extensive assessments of attention to show that different aspects of attention differentially predict language function in aphasia (Murray, 2012). Finally, studies that have explored the role of cognition in aphasia have typically involved a sample of diverse aphasia types and severity. While this is pertinent to capturing the incidence of cognitive impairment in the general aphasic population, the wide variability of aphasia subtypes can confound analyses of the links between domain-general cognitive impairment and any particular aphasic subtype or symptom.

Executive functions and language are closely linked in both brain and behaviour. Behaviourally, cognitive control and working memory have long been known to support language processing (Gordon et al., 2002; Novais-Santos et al., 2007; January et al., 2009; Fedorenko, 2014). Neurally, both executive functions and language robustly engage regions within the left frontal cortex (Kaan and Swaab, 2002; Novick et al., 2005). This makes it challenging to functionally dissociate anatomical correlates of the two domains. Of particular relevance is the function of Broca’s area and the left inferior frontal cortices (LIFC). Damage to Broca’s area, which encompasses cytoarchitecturally defined Brodmann’s area BA 44 and BA 45 of the left posterior inferior frontal gyrus (LpIFG) (Ardila et al., 2016; Papitto et al., 2020) commonly results in anomia, which has led people to believe that Broca’s area within the LpIFG play a causal role in language production. However, research in more recent years challenges this notion. The current view is that long-term speech production outcome following left inferior frontal damage is best explained by a combination of damage to Broca’s area and neighbouring regions including the underlying white matter (Gajardo-Vidal and Lorca-Puls et al., 2021), which was also damaged in Paul Broca’s two historic cases (Dronkers et al., 2007), and that Broca’s area is not specialised for speech and language, but rather is part of a wider network of general cognitive processing that includes, but is not limited to language (Duncan, 2010; Duncan, 2013). Nevertheless, some argue that executive functions and language occupy nearby but distinct regions within the left frontal cortex (Fedorenko and Varley, 2016). To date, the brain areas required for speech production, and the type of aphasia that results from damage to the LIFC remains a topic of continued debate (Marie, 1906; Mohr et al., 1978; Alexander et al., 1990; Lorch, 2008; Fridriksson et al., 2015; Tremblay and Dick, 2016).

Here, we aimed to tease apart the cognitive processes associated with language production, and their underlying neural correlates – with a particular interest in how lesion correlates within the LIFG are associated with long-term language production deficits. We sampled a wide range of PWA from left hemisphere stroke who had long-term speech production deficits (anomia) and varying damage to LIFG. Anomia is the most common symptom of post-stroke aphasia and manifests as difficulty in word retrieval when naming common objects (Laine and Martin, 2013). The participants in this study had relatively intact comprehension and no speech apraxia. The participants were assessed on an extensive battery of language and domain-general cognitive functions. The behavioural data were analysed using Principal Component Analysis (PCA) and their underlying lesion correlates were mapped using Voxel-Based Correlational Morphology (VBCM). We predicted that our analysis would reveal a non-language specific higher-level cognitive component to anomic symptoms, which would be driven by damage to Broca’s area and surrounding regions in the left frontal cortices.

## 2. Materials and methods

### 2.1 Participants

Thirty-six English speakers with chronic aphasia following a single left-hemisphere stroke participated in the study (see Fig. 1 for a lesion overlap map, Table 1 for demographic and clinical data). All were at least 12 months post-stroke and at the time of scanning and assessment, had normal hearing, normal or corrected-to-normal visual acuity and no previous history of significant neurological or psychiatric disease. Inclusion criteria were: (i) anomia as determined by the naming subtest of the Comprehensive Aphasia Test (Swinburn et al., 2005); (ii) good single word comprehension as assessed by the spoken word comprehension subtest of the Comprehensive Aphasia Test (Swinburn et al., 2005); (iii) relatively spared ability to repeat single monosyllabic words from the Psycholinguistic Assessments of Language Processing in Aphasia (Kay et al., 1992); (iv) absence of speech apraxia as determined by the Apraxia Battery for Adults (Dabul, 2000). Participants were excluded if they had any contraindications for scanning or any other significant neurological or psychiatric conditions. Informed consent was obtained from all participants in accordance with the Declaration of Helsinki and the study was approved by the Central London Research Ethics Committee, UK.

**Table 1.** Participant demographic and clinical data

| ID | Gender | Age<br>(years) | Education<br>(years) | Time post-stroke (years) | Total lesion<br>volume (cm<sup>3</sup>) |
| --- | --- | --- | --- | --- | --- |
| 01 | M | 55 | 16 | 11 | 61.53 |
| 02 | M | 56 | 11 | 7 | 160.80 |
| 03 | M | 71 | 13 | 2 | 42.76 |
| 04 | M | 55 | 11 | 9 | 57.24 |
| 05 | M | 71 | 16 | 2 | 78.36 |
| 06 | F | 51 | 13 | 13 | 43.42 |
| 07 | M | 47 | 13 | 12 | 161.81 |
| 08 | F | 66 | 16 | 18 | 83.18 |
| 09 | M | 61 | 16 | 4 | 63.56 |
| 10 | M | 44 | 16 | 3 | 38.14 |
| 11 | F | 44 | 17 | 1 | 29.49 |
| 12 | F | 70 | 11 | 12 | 8.93 |
| 13 | M | 70 | 11 | 29 | 117.55 |
| 14 | M | 69 | 16 | 7 | 171.71 |
| 15 | M | 73 | 11 | 11 | 71.31 |
| 16 | F | 45 | 11 | 3 | 63.53 |
| 17 | F | 53 | 11 | 5 | 22.25 |
| 18 | F | 55 | 13 | 5 | 1.51 |
| 19 | M | 40 | 17 | 8 | 163.75 |
| 20 | M | 64 | 13 | 24 | 308.18 |
| 21 | M | 42 | 17 | 1 | 65.80 |
| 22 | M | 74 | 16 | 11 | 164.53 |
| 23 | M | 63 | 16 | 25 | 156.91 |
| 24 | M | 64 | 16 | 10 | 348.23 |
| 25 | M | 60 | 16 | 11 | 94.32 |
| 26 | F | 60 | 11 | 6 | 223.29 |
| 27 | M | 75 | 11 | 12 | 112.55 |
| 28 | F | 50 | 16 | 3 | 130.60 |
| 29 | M | 64 | 11 | 8 | 240.39 |
| 30 | M | 29 | 13 | 6 | 78.61 |
| 31 | F | 81 | 10 | 16 | 99.38 |
| 32 | M | 60 | 13 | 20 | 403.11 |
| 33 | M | 65 | 13 | 14 | 387.17 |
| 34 | M | 82 | 13 | 34 | 152.61 |
| 35 | M | 58 | 16 | 8 | 239.29 |
| 36 | M | 39 | 17 | 2 | 95.70 |

Thirty-six participants. 10 Female, age range 29-82 years (mean: 59, SD: 12.5). Average time post-stroke was 10 years and average lesion volume was 131.7cm<sup>3</sup>.

**Figure 1.**
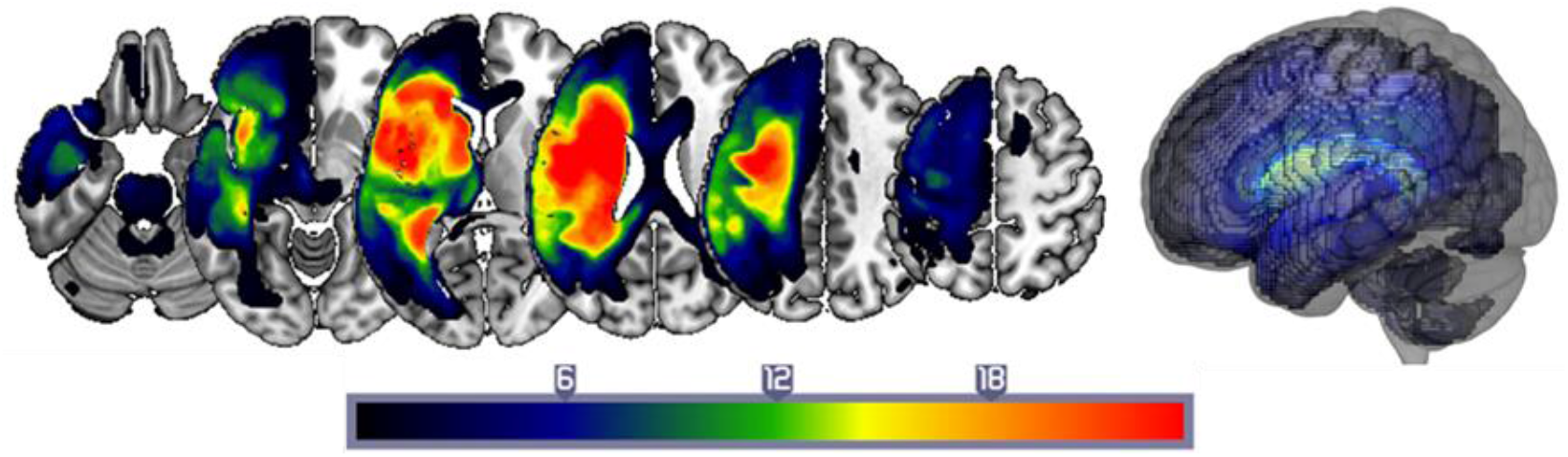
Lesion overlap map. A lesion overlap map for the 36 stroke anomic participants. Colour scale represents frequency of regional brain damage (hot-body scale with red indicating most frequently damaged brain regions i.e., >18 patients, while dark blue < 6 patients with damage to these regions). Results are shown overlaid on the MNI template brain, created in MRIcro-GL (Rorden et al., 2007).

### 2.2 Neuropsychology

#### Behavioural assessment

A comprehensive battery of language and non-language tests was administered to assess participants’ language and cognitive abilities (see supplementary material for all administered behavioural tests [Supplementary Table 1] and percentage of participants with impaired performance scores [supplementary Fig. 1]).

The language tests administered to assess speech production included the naming and repetition subtests of the CAT, the word/non-word repetition subtests from the Psycholinguistic Assessments of Language Processing in Aphasia subtests 8 and 9 (PALPA; Kay et al., 1992), the Boston Naming Test (BNT; Kaplan, Goodglass and Weintraub, 1983). The language assessments that captured other language functions included Pyramids and Palm Trees (PPT; Howard and Patterson, 1992), other subtests from the CAT, and the reading tasks from the PALPA8.

The non-language cognitive assessments included the Cattel Culture Fair IQ Test (Scale 2, Form A; Cattell and Cattell, 1963), Rey-Osterrieth Complex Figure Test (Osterrieth, 1944), Digit Span tasks from the Wechsler Adult Intelligence Scale – Fourth Edition (WAIS-IV; Wechsler, 2008), the trail making and card sorting subtests from the Delis-Kaplan Executive Functions System test (D-KEFS; Delis, Kaplan and Kramer, 2001), the Hopkins Verbal Learning Test (HVLT; Brandt and Benedict, 2001) and the Children’s Sustained Attention to Response Task (cSART; Robertson et al., 1997).

#### Principal Component Analysis (PCA)

PCA is a useful exploratory tool that can extract the underlying latent structure of a set of correlated variables – like scores in standardised assessments of post-stroke cognitive impairment. There has been increasing interest in interpreting these latent variables in terms of the potentially separable cognitive sub-systems underlying (often strongly correlated) task scores. This is typically done by correlating latent variables with the original scores: those scores that correlate more strongly with the latent variable are said to load on that latent variable. Here, following recent results, we employ varimax rotation to encourage greater sparsity, and thus interpretability, in those loadings (Butler et al., 2014; Halai et al., 2017; Tochadse et al., 2018).

Participants’ scores on all assessments were entered into a PCA with varimax rotation (conducted with SPSS 26.0). We had 55 variables and 36 cases. Factors with an eigenvalue ≥1.0 were extracted then rotated. After orthogonal rotation, the factor loadings of each test allowed interpretation of what cognitive-language primary process was represented by that factor (Table 2). Individual participants’ scores on each extracted factor were then used as behavioural covariates in the neuroimaging analysis.

**Table 2.**
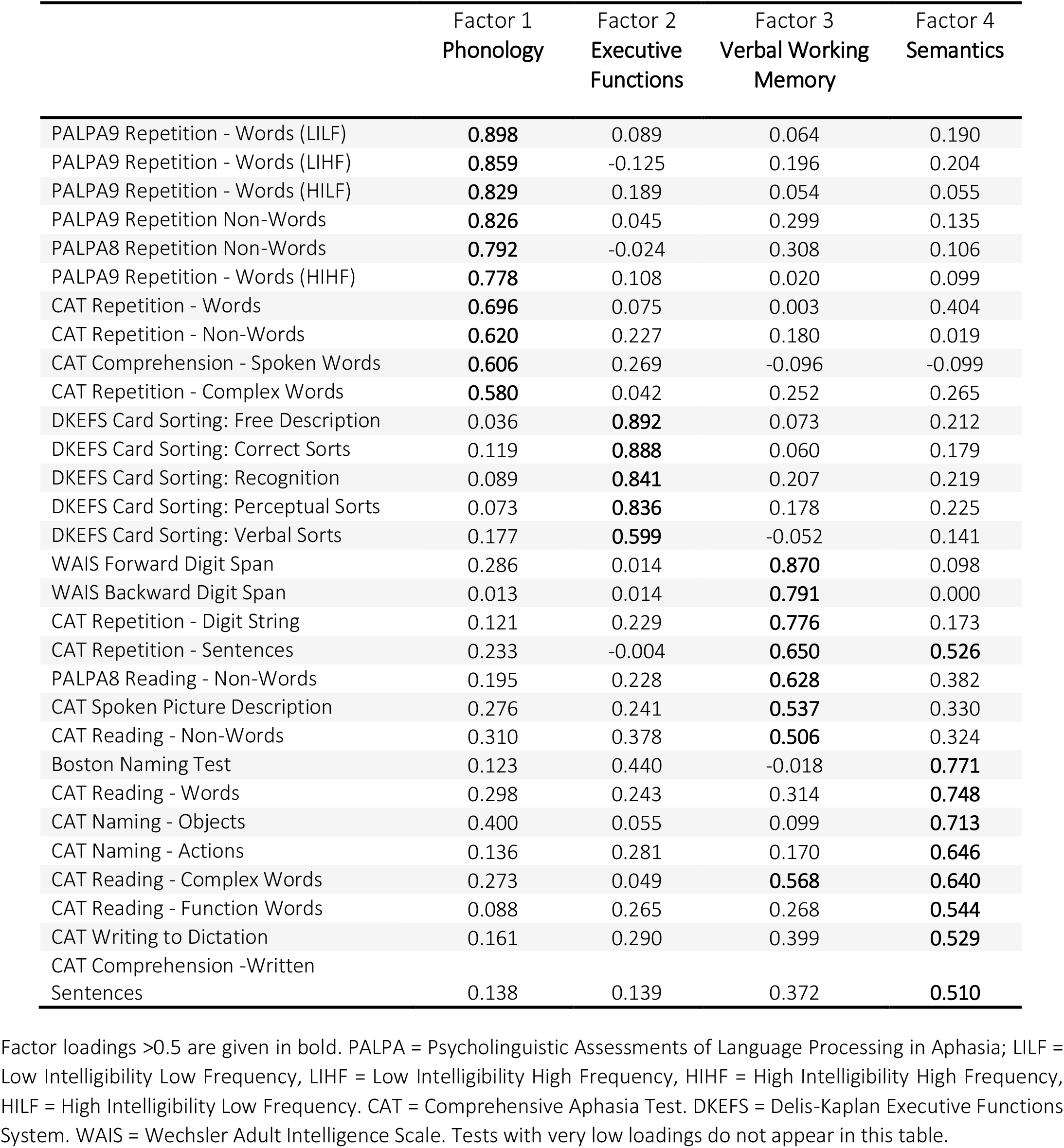
Loadings of behavioural assessments on rotated PCA factors.

### 2.3 Neuroimaging

#### MR Imaging acquisition and analysis

Whole-brain imaging was performed on a 3T Siemens TIM-Trio system (Siemens, Erlangen, Germany) at the Wellcome Centre for Human Neuroimaging. Structural (T1-weighted) MRI images were normalised using Statistical Parametric Mapping software (SPM12) running under Matlab 2015a (MathWorks, Natick, MA). Lesion images were defined by the Automatic Lesion Identification toolbox (ALI; Seghier et al., 2008), employing a variant of the unified segmentation algorithm (Ashburner and Friston, 2005), optimised for use in the focally damaged brain.

Structural MRI scans were pre-processed with Statistical Parametric Mapping software (SPM12: Wellcome Trust Centre for Neuroimaging, http://www.fil.ion.ucl.ac.uk/spm/). The images were normalised into standard Montreal Neurological Institute (MNI) space using a modified unified segmentation–normalisation procedure optimised for focal lesioned brains (Seghier et al., 2008). Data from all participants were entered into the segmentation–normalisation. This procedure combines segmentation, bias correction and spatial normalisation through the inversion of a single unified model (see Ashburner and Friston, 2005 for more details). In brief, the unified model combines tissue class (with an additional tissue class for abnormal voxels), intensity bias and non-linear warping into the same probabilistic models that are assumed to generate subject-specific images. Images were then smoothed with an 8 mm full-width-half-maximum (FWHM) Gaussian kernel and used in the lesion analyses described below. The lesion of each participant was automatically identified using an outlier detection algorithm, compared to healthy controls, based on fuzzy clustering. Voxel values in these regions range from 0 to 1, with higher values indicating greater evidence that the voxel is damaged, and evidence is derived by comparing tissue intensity in each voxel to intensities from a population of neurologically normal controls. The default parameters were used. The images generated for each participant were individually checked and visually inspected with respect to the original scan and were used to create the lesion overlap map in Fig. 1. We selected the Seghier et al. (2008) method as it is objective and efficient for a large sample of lesions (Wilke, de Haan, Juenger and Karnath, 2011).

#### Lesion-Symptom Mapping

For lesion-symptom mapping, we used the fuzzy lesion images as described above and correlated these with PCA factor scores using a voxel-based correlational methodology (VBCM: Tyler, Marslen-Wilson and Stamatakis, 2005), a variant of voxel-lesion symptom mapping (VLSM: Bates et al., 2003). We used VBCM because this approach i) has the virtue of preserving the continuous nature of both behavioural and neural indices i.e., does not require a binary classification of the intact/lesioned brain to be marked, as in the case of VLSM, and ii) replicates previous methodology using varimax-rotated PCA in aphasia (e.g. Butler et al., 2014), aiding data comparisons within the field.

The VBCM analysis of PCA factors was conducted in SPM12 running on Matlab 2019b. The analysis used the four continuous multidimensional predictors of the PCA factor scores, which are necessarily uncorrelated (orthogonal) with one another; these were entered simultaneously as continuous behavioural covariates. The outcome of the analysis therefore denotes which voxels’ variation in tissue concentration corresponds to the unique variance in a given principal component, while controlling for variation in the other components in the analysis. In order to ensure that the results were not merely attributable to lesion size, each participants’ lesion volume was calculated from the lesion identified by the automated lesion identification method (Seghier et al., 2008) and this was entered as a covariate in the VBCM. All analyses were performed with and without a correction for lesion volume. All anatomical labels were based on the Harvard–Oxford atlas in MNI space.

## 3. Results

### 3.1 Neuropsychological profiles and principal language-cognitive factors

The rotated PCA produced a four-factor solution which accounted for 55% of variance in participants’ performance (F1 = 28.6%; F2 = 10.6%; F3 = 8.3%; F4 = 7.1%). The loadings of each of the different behavioural assessments on each of the factors are given in Table 2 (for individual participants’ scores on each factor and percentage of participants with impaired language and non-language scores, see supplementary Table 1 and supplementary Fig. 2 respectively). Tasks that tapped into input and output phonology (e.g. word and non-word repetition) loaded heavily on Factor 1, as such we refer to this factor as ‘Phonology’. Factor 2 was interpreted as ‘Executive Functions’, as assessments that loaded most heavily on it tapped into non-verbal cognitive processes (e.g. problem solving and concept formation). Assessments that loaded on Factor 3 were those requiring speech output (e.g., composite picture description) and online maintenance and use of auditory inputs (e.g. digit span, sentence repetition) along with phonological skills (e.g. reading aloud non-words), hence we refer to this factor as ‘verbal working memory’. Finally, Factor 4 was interpreted as ‘Semantics’, the assessments that loaded on this factor were more diverse but primarily required processing of meaning (e.g. picture naming and comprehension of written sentences).

### 3.2 The neural basis of performance in chronic stroke aphasia

#### Voxel-based morphometry of principal component analysis factors

The VBCM results are shown in Fig. 2 and Table 3. Each map displays where tissue damage covaries uniquely with a given factor score, where the factors are necessarily uncorrelated with one another. Results are thresholded at p ≤ 0.001 voxel-level and p < 0.05 FWE corrected at cluster-level.

**Figure 2.**
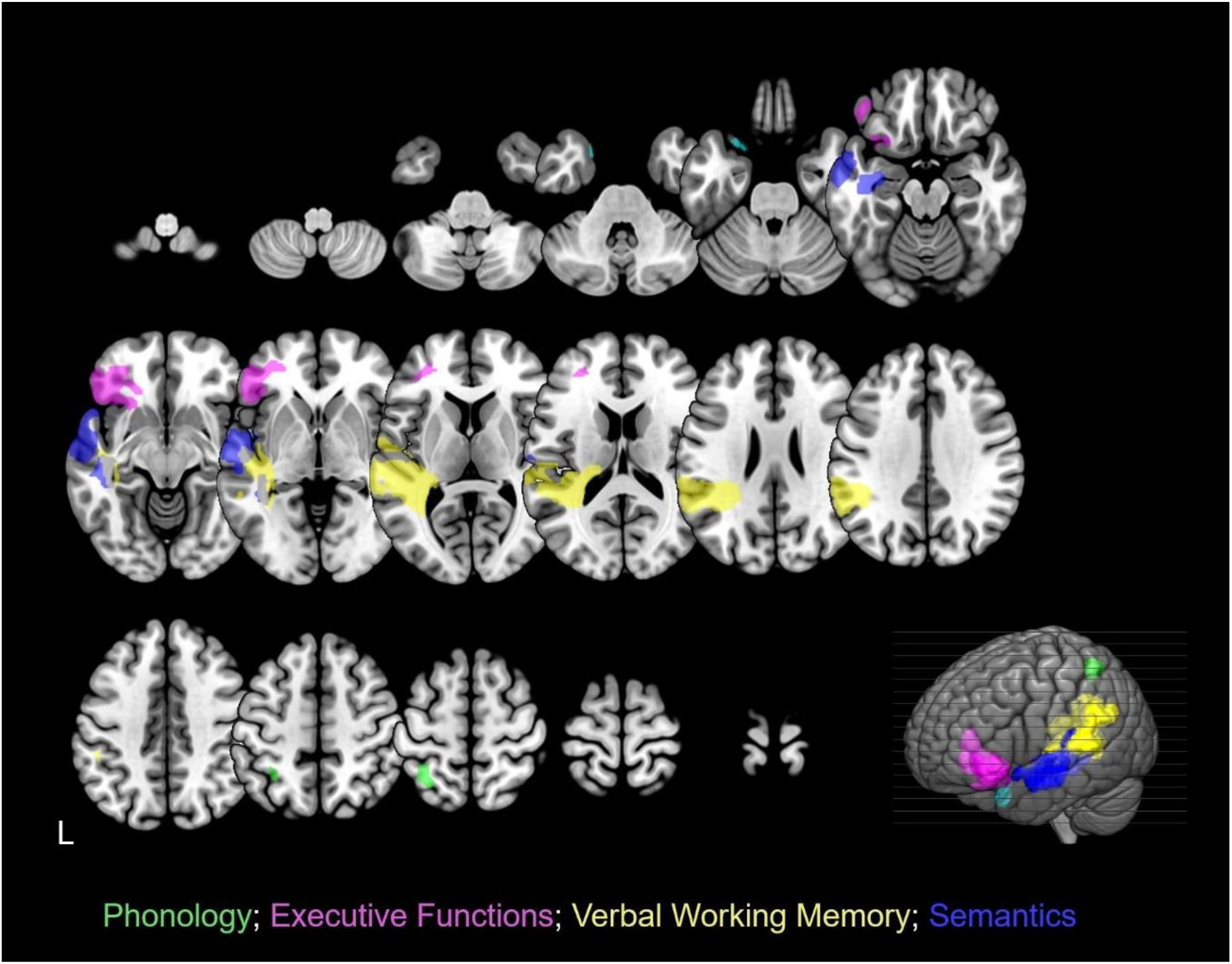
Structural correlates associated with each component from the combined PCA. Phonology: green; Executive Functions: magenta; Verbal Working Memory: yellow; Semantics: two distinct clusters in cyan and indigo. Clusters were obtained by applying a voxel-level threshold at p ≤ 0.001 and a family-wise error correction of p < 0.05 at cluster level. The lower right corner displays a rendered template brain (created in MRIcro-GL) showing the significant clusters projected to the left brain surface.

**Table 3.** Neural correlates for omnibus PCA factors.

| Principal Component | Location | Extent (voxels) | Z | MNI co-ordinates |  |  |
| --- | --- | --- | --- | --- | --- | --- |
|  |  |  |  | x | y | z |
| F1 (Phonology) | Left Superior Parietal Lobe | 175 | 4.16 | -34 | -54 | 58 |
| F2 (Executive Functions) |  | 1563 |  |  |  |  |
|  | Left Inferior Frontal Gyrus (Pars Triangularis) |  | 3.93 | -44 | 36 | 2 |
|  | Left Inferior Frontal Gyrus (Pars Orbitalis) |  | 3.59 | -40 | 42 | -8 |
|  | Left Middle Frontal Gyrus (Dorsolateral Prefrontal Cortex) |  | 3.56 | -30 | 44 | 4 |
| F3 (Verbal Working Memory) |  | 5262 |  |  |  |  |
|  | Left Posterior Superior Temporal Gyrus |  | 4.49 | -58 | -28 | 6 |
|  | Left Superior Longitudinal Fasciculus |  | 4.94 | -38 | -44 | 18 |
|  | Left Posterior Thalamic Radiation |  | 5.35 | -36 | -46 | 2 |
| F4 (Semantics) |  | 209 |  |  |  |  |
|  | Left Superior Temporal Pole |  | 4.82 | -22 | 10 | -24 |
|  | Left Middle Temporal Pole |  | 4.47 | -20 | 12 | -34 |
|  |  | 2343 |  |  |  |  |
|  | Left Superior Temporal Gyrus |  | 4.14 | -64 | -10 | -4 |
|  | Left Middle Temporal Gyrus |  | 4.14 | -62 | -14 | -12 |

Only clusters with cluster-level FWEc $p < 0.05$ are shown in the table.

Performance on the phonological factor was uniquely correlated with a cluster of voxels in the left parietal lobe, with peak voxels in the left superior parietal lobe. The cluster also included voxels in the left inferior parietal lobule.

Performance on the executive functions factor was uniquely related to a cluster of voxels in the left frontal lobe, with peak voxels in the left inferior frontal gyrus (pars orbitalis and pars triangularis) and the left dorsolateral prefrontal cortex.

Performance on the verbal working memory factor was uniquely related to a large cluster of voxels in the left hemisphere, with peak voxels in the posterior superior temporal gyrus, the superior longitudinal fasciculus and posterior thalamic radiation. The cluster also included voxels within left Werenicke’s area, Heschl’s gyrus and the hippocampus.

Performance on the semantic factor was uniquely related to two clusters in the left hemisphere with peak voxels in the superior/ middle temporal pole and superior/middle temporal gyrus. The clusters also included voxels across the left insula.

#### Lesion size and age

Given that some brain regions are more likely than others to be damaged after middle cerebral artery (MCA) stroke (Phan et al., 2005) and some regions are more susceptible to age-related atrophy, we controlled for lesion volume and age in subsequent lesion-symptom analyses.

Each participant’s lesion volume was calculated from the lesion identified by the modified segmentation-normalization procedure (see ‘Materials and methods’ section). For the PCA factors, lesion volume correlated relatively weakly with the phonology factor (r=0.137, p=0.426), the auditory working memory factor (r= -0.318, p=0.059) and semantic factor (r= -0.313, p=0.063), and slightly more strongly with the executive-functions factor (r= -0.426, p=0.10)

Including age in the VBCM model with the PCA factor scores did not alter the pattern of results obtained. However, including lesion volume in the model reduced the significance of the executive functions measure, which only reached suprathreshold at voxel-level p < 0.05 and FWEc cluster-level p < 0.05, but did not alter the pattern of results in the remaining 3 PCA factors. As previously mentioned, the executive functions component correlated with tissue damage in the left inferior frontal cortex (LIFC); as a common region of damage following left MCA stroke (Phan et al., 2005), high covariance between LIFC tissue integrity and total lesion volume is expected.

## 4. Discussion

The aim of the current study was to investigate the presence of latent cognitive factors that might explain the variance in aphasic language production abilities and how this relates to underlying lesion patterns. We conducted an extensive language and non-language neuropsychological assessment in a sample of thirty-six PWA with long-term language production deficits. Our results replicate and extend work on the neural correlates of higher-level cognitive functions in PWA and their role in language production. We show that (i) the variance underlying language and non-language test performance was best captured by four orthogonal components, two higher-order cognitive components (executive functions and verbal working memory) and two linguistic processing components (phonology and semantics) (Table 2); (ii) brain-behaviour relationships revealed separable neural correlates for each component in line with previous studies and showed that lesions to the left inferior frontal cortex (LIFC) are associated with executive dysfunction, independent of language ability (Fig. 2, Table 3), suggesting that these regions are involved in, but not specific to, language production.

The neural correlates associated with the two language components were in line with previous literature. The phonological component explained the largest proportion of behavioural variance in our group of anomic adults. Scores on this component, which in our study loaded principally on tests of single word and non-word repetition, uniquely correlated with tissue damage in the left superior and inferior parietal lobule (Fig. 2). This is in line with work showing impaired speech repetition following left hemisphere stroke is associated with left parietal lobe damage (Fridriksson et al., 2010). More recent studies that have used a similar approach to ours, with a combined rotated PCA and VBCM in people with aphasia reported a phonology component uniquely related to left temporo-parietal regions (Butler et al.,2014; Halai et al., 2017; Schumacher et al., 2019; Alyahya et al., 2020). It is important to note that the phonology component in those studies also loaded on tests of naming and verbal working memory, as well as repetition, whereas our phonology component was specific to input/output phonology and loaded heavily on tests of single word and non-word repetition. The semantic component explained the least amount of behavioural variance in our sample. Scores on this factor loaded on tests of naming, reading and written comprehension and uniquely correlated with regions in the left superior/ medial temporal pole and the left superior/ medial temporal gyrus (Fig. 2). This supports recent findings that extend the temporal regions implicated in semantic processing (Jackson, 2021).

Importantly, higher cognitive functions, namely executive functions and verbal working memory, independently explain a significant amount of variance in language abilities in our population of aphasics with chronic language production deficits. Both have also been shown to be robust behavioural predictors of aphasia recovery outcomes (Fillingham, Sage and Lambon Ralph, 2005a, 2005b, 2006; Conroy, Sage and Lambon Ralph, 2009; Lambon Ralph et al., 2010; Yeung and Law, 2010; Snell, Sage and Lambon Ralph, 2010, Sage, Snell and Lambon Ralph, 2011; Dignam et al., 2017). During aphasia recovery, executive functions are argued to be important for the generation of semantic and phonological concepts to aid with word retrieval (Dignam et al, 2017) and to navigate other complex dynamics of human communication, while the integrity of general memory processes enables (re)learning and retention of linguistic knowledge during rehabilitation. Schumacher and colleagues (2020) show that variance in functional communication abilities in PWA, as measured by the Amsterdam Nijmegen Everyday Language Test, can be almost entirely accounted for by patients’ verbal short-term memory. In our study, the verbal working memory component uniquely correlated with regions of tissue damage in the left posterior superior temporal gyrus, left superior longitudinal fasciculus, as well as Heschl’s gyrus, Wernicke’s area and the hippocampus (Fig. 2). This component captured abilities both in continuous (narrative) speech production (e.g, spoken picture description) and online maintenance of increasing auditory information (e.g. digit-span, sentence repetition). This replicates findings from Tochadse et al., (2019) who report a similar neural correlate associated with auditory working memory in PWA.

Scores on the executive functions factor uniquely correlated with tissue damage in the left inferior frontal cortex (LIFC), including pars orbitalis and pars triangularis, and middle frontal gyrus (DLPFC) (see Fig. 2 for structural correlates and table 3 for MNI co-ordinates). The LIFC results support and extends recent findings from ECoG and fMRI. Conner et al., (2019) used intracranial recordings to show that activity in pars triangularis and pars orbitalis is specifically engaged in object naming, compared to scrambled images, and shows stronger activity for words with high selectivity (number of possible correct responses). Ekert et al., (2021) used fMRI to show that pars orbitalis was most activated during object naming, compared to repetition of words and pseudowords. Our participants all had anomia, and by definition significant object naming deficits. However, our results show that lesions to pars triangularis and pars orbitalis are associated with executive functions, independent of language function. This suggests that these regions within the LIFC support high-level planning and execution that is important for object naming, but not specific to language processing. This supports contemporary models of speech and language that suggest that language production may rely on the same process and neural systems that support other high-level action planning and execution (Botvinivk, 2008; Hickok 2012; Weiss et al., 2016). These findings support the role of Broca’s area, here pars triangularis in particular, and adjacent pars orbitalis in domain-general cognition and extend our understanding of the neural correlates of anomia. We show that in a group of PWA with chronic anomia, lesions to the LIFC including Broca’s area are associated with executive (dys)function, independent of language abilities. This suggests that, while damage to the LIFC commonly coincides with language impairment after stroke, lesions to this area might be driving a (non-language specific) cognitive component of anomia that co-occurs with language impairment We speculate that lesions to Broca’s area may lead to deficits in high-level executive functioning that supports language production and that this can contribute to varying levels of long-term language impairment the nature of which will vary depending on the pattern of damage to neighbouring regions of grey and white matter (Kimberg et al., 2007; Richardson et al., 2012; Inoue et al., 2014; Mah et al., 2014; Sperber and Karnath et al., 2017; Gajardo-Vidal and Lorca-Puls et al., 2021).

Behaviourally, the executive functions component loaded on tests of problem solving and concept formation as measured by the D-KEFS Card Sorting assessment. Card sorting assessments, including the D-KEFS and Wisconsin (Berg, 1948) tasks, appear to reliably engage executive functions and relate to damage in the left inferior frontal cortices (LIFC) and DLPFC in our group of aphasic adults. The neural correlates associated with our executive functions component show some overlap, namely pars triangularis and DLPFC, with a PCA component identified by Schumacher and colleagues (2019), which the authors refer to as ‘inhibit-generate’. Their ‘inhibit-generate’ component captured abilities of idea generation, reasoning, problem solving and response inhibition in PWA and loaded on, amongst others, the D-KEFS card sorting test, as we used here. DLPFC is also reported by Lacey et al., (2017) as a neural correlate of their executive functions component which loaded on, amongst others, tests of planning, rule following and cognitive flexibility in PWA. Alyahya and colleagues (2020) also identify the middle frontal gyrus as a structural correlate of executive functions, specifically tests of abstract reasoning and rule following, in aphasic adults. Baldo et al. (2005) reported impairment on the Wisconsin Card Sorting Task in aphasic individuals, but not in adults with left-hemisphere brain damage without aphasia, suggesting that the card sorting task taps into executive functions that are necessary for effective language function. Consistent with this, Dignam et al., (2017) show that the D-KEFS Card Sorting assessment is predictive of successful anomia therapy outcomes. Collectively, these findings suggest that in PWA, card sorting tasks such as the D-KEFS, that we used here, are a sensitive measure of executive functions supporting language functioning. Not including these assessments of concept formation and problem solving skills might be a significant contributing reason to why previous studies in aphasia have previously struggled to find consistent associations between tests of executive functions and brain damage (Kummerer et al., 2013; Butler et al., 2014; Mirman et al., 2015; Halai et al., 2017; Tochadse et al., 2018).

In conclusion, our findings suggest that in people with chronic post-stroke anomia, cognitive abilities and in particular executive functions and verbal working memory, help explain significant variance in language function, more than classical purely linguistic models of the condition imply. Moreover, lesions to the LIFC, including Broca’s area, determine whether people suffer worse executive (dys)function, independent of their language abilities. This does not necessarily imply that all aphasics will have additional cognitive impairments, but that in those who do, higher-level executive functions may explain more of the variance in language production ability than previously thought. A better understanding of the covariance between language and non-language deficits and their underlying neural correlates will inform more targeted aphasia treatment, tailored to an individual’s pattern of impairments. This may be in the form of neurostimulation targeting regions of domain-general cognition or by incorporating measures of higher-order cognitive function, such as concept formation and verbal working memory, to improve the accuracy of aphasia prediction models (Price et al., 2010; Hope et al., 2013, 2018; Yourganov et al., 2015).

## References

1. Alexander, M. P., Naeser, M. A. and Palumbo, C. (1990) ‘ Brocas area aphasias - aphasia after lesions including the frontal operculum’, Neurology,40 (2), pp.353–362.

2. Alyahya, R. S. W., Halai, A. D., Conroy, P., Lambon Ralph, M.A. (2020), ‘A unified model of post-stroke language deficits including discourse production and their neural correlates’, Brain, 143(5), pp.1541–1554.

3. Ardila, A., Bernal, B. and Rosselli, M. (2016) ‘How Localized are Language Brain Areas? A Review of Brodmann Areas Involvement in Oral Language’, Archives of Clinical Neuropsychology, 31(1), pp. 112–122.

4. Ashburner, J. and Friston, K. J. (2005) ‘Unified segmentation’, Neuroimage, 26(3), pp. 839–851.

5. Baldo, J., Dronkers, N., Wilkins, D., Ludy, C., Raskin, P. and Kim, J. (2005) ‘Is problem solving dependent on language?’, Brain and Language, 92(3), pp. 240–250.

6. Bates, E., Wilson, S. M., Saygin, A. P., Dick, F., Sereno, M. I., Knight, R. T. and Dronkers, N. F. (2003) ‘Voxel-based lesion–symptom mapping’, Nature Neuroscience, 6(5), pp. 448–450.

7. Berg, E. A. (1948) ‘A Simple Objective Technique for Measuring Flexibility in Thinking’, The Journal of General Psychology, 39(1), pp. 15–22.

8. Botvinick, M. M. (2008), ‘Hierarchical models of behaviour and prefrontal function’, Trends in Cognitive Sciences, 12, pp. 201–208.

9. Brandt J, Benedict R. Verbal Learning Test-Revised Professional Manual. Lutz, FL: Psychological Assessment Resources, Inc (2001).

10. Butler, R. A., Lambon Ralph, M.A. and Woollams, A. M. (2014) ‘Capturing multidimensionality in stroke aphasia: mapping principal behavioural components to neural structures’, Brain, 137(12), pp. 3248–3266.

11. Cattell, R. B. and Cattell, A. K. S. (1963). Culture fair intelligence test. Champaign, IL: Institute for Personality and Ability Testing

12. Conner, C. R., Kadipasaoglu, C. M., Shouval. H.Z., Hickok, G., Tandon, N. (2019), ‘Network dynamics of Broca’s area during word selection’. PLoS ONE 14(12): e0225756.

13. Conroy, P., Sage, K. and Lambon Ralph, M.A. (2009) ‘The effects of decreasing and increasing cue therapy on improving naming speed and accuracy for verbs and nouns in aphasia’, Aphasiology, 23(6), pp. 707–730.

14. Dabul, B. (2000). Apraxia battery for adults (second ed.). Austin, Tx: Pro-Ed.

15. Delis, D. C., Kaplan, E., and Kramer, J. H. (2001). D-KEFS Executive Function System: Examiners manual. San Antonio, TX: Psychological Corporation.

16. Dignam, J., Copland, D., O’Brien, K., Burfein, P., Khan, A. and Rodriguez, A. D. (2017) ‘Influence of Cognitive Ability on Therapy Outcomes for Anomia in Adults With Chronic Poststroke Aphasia’, Journal of Speech, Language, and Hearing Research, 60(2), pp. 406–421.

17. Dronkers, N. F., Plaisant, O., Iba-Zizen, M. T. and Cabanis, E. A. (2007) ‘Paul Broca’s historic cases: high resolution MR imaging of the brains of Leborgne and Lelong’, Brain, 130(5), pp. 1432–1441.

18. Duncan, J. (2010) ‘The multiple-demand (MD) system of the primate brain: mental programs for intelligent behaviour’, Trends in Cognitive Sciences, 14(4), pp. 172–179.

19. Duncan, J. (2013) ‘The Structure of Cognition: Attentional Episodes in Mind and Brain’, Neuron, 80(1), pp. 35–50.

20. Ekert, J. O., Lorca-Puls, D. L., Gajardo-Vidal, A., Crinion, J. T., Hope, T. M. H., Green, D. W., Price, C. J. (2021), ‘A functional dissociation of the left frontal regions that contribute to single word production tasks’, Neuroimage, 245, pp. 118734.

21. El Hachioui, H., Visch-Brink, E. G., Lingsma, H. F., Van De Sandt-Koenderman, M. W. M. E., Dippel, D. W. J., Koudstaal, P. J. and Middelkoop, H. A. M. (2014) ‘Nonlinguistic Cognitive Impairment in Poststroke Aphasia’, Neurorehabilitation and Neural Repair, 28(3), pp. 273–281.

22. Fedorenko, E. (2014) ‘The role of domain-general cognitive control in language comprehension’, Frontiers in Psychology, 5.

23. Fedorenko, E. and Varley, R. (2016) ‘Language and thought are not the same thing: evidence from neuroimaging and neurological patients’, Annals of the New York Academy of Sciences, 1369(1), pp. 132–153.

24. Fillingham, J., Sage, K. and Lambon Ralph, M. (2005a) ‘Further explorations and an overview of errorless and errorful therapy for aphasic word-finding difficulties: The number of naming attempts during therapy affects outcome’, Aphasiology, 19(7), pp. 597–614.

25. Fillingham, J. K., Sage, K. and Lambon Ralph, M.A. (2005b) ‘Treatment of anomia using errorless versus errorful learning: are frontal executive skills and feedback important?’, International Journal of Language & Communication Disorders, 40(4), pp. 505–523.

26. Fillingham, J. K., Sage, K. and Lambon Ralph †, M.A. (2006) ‘The treatment of anomia using errorless learning’, Neuropsychological Rehabilitation, 16(2), pp. 129–154.

27. Frankel, T., Penn, C. and Ormond-Brown, D. (2007) ‘Executive dysfunction as an explanatory basis for conversation symptoms of aphasia: A pilot study’, Aphasiology, 21(6-8), pp. 814–828.

28. Fridriksson, J., Fillmore, P., Guo, D. and Rorden, C. (2015) ‘Chronic Broca’s Aphasia Is Caused by Damage to Broca’s and Wernicke’s Areas’, Cerebral Cortex, 25(12), pp. 4689–4696.

29. Fridriksson, J., Kjartansson, O., Morgan, P. S., Hjaltason, H., Magnusdottir, S., Bonilha, L. and Rorden, C. (2010) ‘Impaired Speech Repetition and Left Parietal Lobe Damage’, Journal of Neuroscience, 30(33), pp. 11057–11061.

30. Fridriksson, J., Nettles, C., Davis, M., Morrow, L. and Montgomery, A. (2006) ‘Functional communication and executive function in aphasia’, Clinical Linguistics & Phonetics, 20(6), pp. 401–410.

31. Fucetola, R., Connor, L. T., Strube, M. J. and Corbetta, M. (2009) ‘Unravelling nonverbal cognitive performance in acquired aphasia’, Aphasiology, 23(12), pp. 1418–1426.

32. Gajardo-Vidal, A., Lorca-Puls, D. L., Team, P., Warner, H., Pshdary, B., Crinion, J. T., Leff, P., Hope, T. M. H., Geva, S., Seghier, M. L., Green, D. W., Bowman, H. and Price, C. J. (2021) ‘Damage to Broca’s area does not contribute to long-term speech production outcome after stroke’, Brain.

33. Goodglass, H., Kaplan, E., and Barresi, B. (2001) The assessment of aphasia and related disorders (3rd ed.). Philadelphia: Lippincott, Williams, & Wilkins.

34. Gordon, P. C., Hendrick, R. and Levine, W. H. (2002) ‘Memory-Load Interference in Syntactic Processing’, Psychological Science, 13(5), pp. 425–430.

35. Halai, A. D., Woollams, A. M. and Lambon Ralph, M. A. (2017) ‘Using principal component analysis to capture individual differences within a unified neuropsychological model of chronic post-stroke aphasia: Revealing the unique neural correlates of speech fluency, phonology and semantics’, Cortex, 86, pp. 275–289.

36. Helm-Estabrooks, N. (2002) ‘Cognition and aphasia: a discussion and a study’, Journal of Communication Disorders, 35(2), pp. 171–186.

37. Hickok, G. S. (2012), ‘Computational neuroanatomy of speech production’, Nature Reviews Neuroscience, 13, pp. 135–145.

38. Hope, T. M. H., Leff, A. P. and Price, C. J. (2018) ‘Predicting language outcomes after stroke: Is structural disconnection a useful predictor?’, NeuroImage: Clinical, 19, pp. 22–29.

39. Hope, T. M. H., Seghier, M. L., Leff, A. P. and Price, C. J. (2013) ‘Predicting outcome and recovery after stroke with lesions extracted from MRI images’, NeuroImage: Clinical, 2, pp. 424–433.

40. Howard, D., and Patterson, K. (1992) Pyramids and Palm Trees: A test of semantic access from pictures and words. Bury St. Edmunds, UK: Thames Valley Test Company.

41. Inoue, K., Madhyastha, T., Rudrauf, D., Mehta, S. and Grabowski, T. (2014) ‘What affects detectability of lesion–deficit relationships in lesion studies?’, NeuroImage: Clinical, 6, pp. 388–397.

42. Jackson, R. L. (2021) ‘The neural correlates of semantic control revisited’, NeuroImage, 224, pp. 117444.

43. January, D., Trueswell, J. C. and Thompson-Schill, S. L. (2009) ‘Co-localization of Stroop and Syntactic Ambiguity Resolution in Broca’s Area: Implications for the Neural Basis of Sentence Processing’, Journal of Cognitive Neuroscience, 21(12), pp. 2434–2444.

44. Jefferies, E., Patterson, K. and Ralph, M. A. L. (2008) ‘Deficits of knowledge versus executive control in semantic cognition: Insights from cued naming’, Neuropsychologia, 46(2), pp. 649–658.

45. Kaan, E. and Swaab, T. Y. (2002) ‘The brain circuitry of syntactic comprehension’, Trends in Cognitive Sciences, 6(8), pp. 350–356.

46. Kaplan, E., Goodglass, H., Weintraub, S (1983) Boston Naming Test. Philadelphia: Lea & Febiger.

47. Kay, J., Lesser, R., & Coltheart, M. (1992) Psycholinguistic Assessments of Language Processing in Aphasia (PALPA). Hove: Erlbaum.

48. Kimberg, D. Y., Coslett, H. B. and Schwartz, M. F. (2007) ‘Power in Voxel-based Lesion-Symptom Mapping’, Journal of Cognitive Neuroscience, 19(7), pp. 1067–1080.

49. Kümmerer, D., Hartwigsen, G., Kellmeyer, P., Glauche, V., Mader, I., Klöppel, S., Suchan, J., Karnath, H.-O., Weiller, C. and Saur, D. (2013) ‘Damage to ventral and dorsal language pathways in acute aphasia’, Brain, 136(2), pp. 619–629.

50. Lacey, E. H., Skipper-Kallal, L. M., Xing, S., Fama, M. E. and Turkeltaub, P. E. (2017) ‘Mapping Common Aphasia Assessments to Underlying Cognitive Processes and Their Neural Substrates’, Neurorehabilitation and Neural Repair, 31(5), pp. 442–450.

51. Laine, M. and Martin, N. (2013) Anomia: Theoretical and clinical aspects. Psychology Press.

52. Lambon Ralph, M.A., Snell, C., Fillingham, J. K., Conroy, P. and Sage, K. (2010) ‘Predicting the outcome of anomia therapy for people with aphasia post CVA: Both language and cognitive status are key predictors’, Neuropsychological Rehabilitation, 20(2), pp. 289–305.

53. Lee, B. and Pyun, S.-B. (2014) ‘Characteristics of Cognitive Impairment in Patients With Post-stroke Aphasia’, Annals of Rehabilitation Medicine, 38(6), pp. 759.

54. Lorch, M. P. (2008) ‘The merest Logomachy: The 1868 Norwich discussion of aphasia by Hughlings Jackson and Broca’, Brain, 131, pp. 1658–1670.

55. Mah, Y.-H., Husain, M., Rees, G. and Nachev, P. (2014) ‘Human brain lesion-deficit inference remapped’, Brain, 137(9), pp. 2522–2531.

56. Marie P. (1906) Aphasia from 1861 to 1866. Essay of historical criticism on the genesis of the doctrine of aphasia. Sem Méd. 26:565–571.

57. Marinelli, C. V., Spaccavento, S., Craca, A., Marangolo, P. and Angelelli, P. (2017) ‘Different Cognitive Profiles of Patients with Severe Aphasia’, Behavioural Neurology, 2017, pp. 15.

58. Mirman, D., Chen, Q., Zhang, Y., Wang, Z., Faseyitan, O. K., Coslett, H. B. and Schwartz, M. F. (2015) ‘Neural organization of spoken language revealed by lesion–symptom mapping’, Nature Communications, 6(1), pp. 6762.

59. Mohr, J. P., Pessin, M. S., Finkelstein, S., Funkenstein, H. H., Duncan, G. W. and Davis, K. R. (1978) ‘BROCA APHASIA - PATHOLOGIC AND CLINICAL’, Neurology, 28(4), pp. 311–324.

60. Murray, L. L. (2012) ‘Attention and Other Cognitive Deficits in Aphasia: Presence and Relation to Language and Communication Measures’, American Journal of Speech-Language Pathology, 21(2), pp. S51–S64.

61. Nicholas, L. E., Brookshire, R. H., Maclennan, D. L., Schumacher, J. G. and Porrazzo, S. A. (1989) ‘Revised administration and scoring procedures for the Boston Naming test and norms for non-brain-damaged adults’, Aphasiology, 3(6), pp. 569–580.

62. Novais-Santos, S., Gee, J., Shah, M., Troiani, V., Work, M. and Grossman, M. (2007) ‘Resolving sentence ambiguity with planning and working memory resources: Evidence from fMRI’, NeuroImage, 37(1), pp. 361–378.

63. Novick, J. M., Trueswell, J. C. and Thompson-Schill, S. L. (2005) ‘Cognitive control and parsing: Reexamining the role of Broca’s area in sentence comprehension’, Cognitive, Affective, & Behavioral Neuroscience, 5(3), pp. 263–281.

64. Osterrieth, P.A. (1944) Le test de copie d’une figure complexe. Arch. Psychol. 30, 206–356.

65. Papitto, G., Friederici, A. D. and Zaccarella, E. (2020) ‘The topographical organization of motor processing: An ALE meta-analysis on six action domains and the relevance of Broca’s region’, NeuroImage, 206, pp. 116321.

66. Phan, T. G., Donnan, G. A., Wright, P. M. and Reutens, D. C. (2005) ‘A Digital Map of Middle Cerebral Artery Infarcts Associated With Middle Cerebral Artery Trunk and Branch Occlusion’, Stroke, 36(5), pp. 986–991.

67. Price, C. J., Seghier, M. L. and Leff, A. P. (2010) ‘Predicting language outcome and recovery after stroke: the PLORAS system’, Nature Reviews Neurology, 6(4), pp. 202–210.

68. Ramsey, L. E., Siegel, J. S., Lang, C. E., Strube, M., Shulman, G. L. and Corbetta, M. (2017) ‘Behavioural clusters and predictors of performance during recovery from stroke’, Nature Human Behaviour, 1(3), pp. 0038.

69. Richardson, J. D., Fillmore, P., Rorden, C., Lapointe, L. L. and Fridriksson, J. (2012) ‘Re-establishing Broca’s initial findings’, Brain and Language, 123(2), pp. 125–130.

70. Robertson, I. H., Manly, T., Andrade, J., Baddeley, B. T. and Yiend, J. (1997) ‘`Oops!’: Performance correlates of everyday attentional failures in traumatic brain injured and normal subjects’, Neuropsychologia, 35(6), pp. 747–758.

71. Sage, K., Snell, C. and Lambon Ralph, M.A. (2011) ‘How intensive does anomia therapy for people with aphasia need to be?’, Neuropsychological Rehabilitation, 21(1), pp. 26–41.

72. Schumacher, R., Halai, A. D. and Lambon Ralph, M.A. (2019) ‘Assessing and mapping language, attention and executive multidimensional deficits in stroke aphasia’, Brain, 142(10), pp. 3202–3216.

73. Schumacher, R., Bruehl, S., Halai, A. D., & Lambon Ralph, M.A. (2020), ‘The verbal, non-verbal and structural bases of functional communication abilities in aphasia’, Brain communications, 2(2), fcaa118.

74. Seghier, M. L., Ramlackhansingh, A., Crinion, J., Leff, A. P. and Price, C. J. (2008) ‘Lesion identification using unified segmentation-normalisation models and fuzzy clustering’, Neuroimage, 41(4), pp. 1253–1266.

75. Snell, C., Sage, K. and Lambon Ralph, M.A. (2010) ‘How many words should we provide in anomia therapy? A meta-analysis and a case series study’, Aphasiology, 24(9), pp. 1064–1094.

76. Sperber, C. and Karnath, H.-O. (2017) ‘Impact of correction factors in human brain lesion-behavior inference’, Human Brain Mapping, 38(3), pp. 1692–1701.

77. Swinburn K, Porter G, Howard D. (2005) The Comprehensive Aphasia Test. Hove, UK: Psychology Press.

78. Tochadse, M., Halai, A. D., Ralph, M. A. L. and Abel, S. (2018) ‘Unification of behavioural, computational and neural accounts of word production errors in post-stroke aphasia’, Neuroimage-Clinical, 18, pp. 952–962.

79. Tranter, L. J. and Koutstaal, W. (2008) ‘Age and Flexible Thinking: An Experimental Demonstration of the Beneficial Effects of Increased Cognitively Stimulating Activity on Fluid Intelligence in Healthy Older Adults’, Aging, Neuropsychology, and Cognition, 15(2), pp. 184–207.

80. Tremblay, P. and Dick, A. S. (2016) ‘Broca and Wernicke are dead, or moving past the classic model of language neurobiology’, Brain and Language, 162, pp. 60–71.

81. Tyler, L. K., Marslen-Wilson, W. and Stamatakis, E. A. (2005) ‘Dissociating neuro-cognitive component processes: voxel-based correlational methodology’, Neuropsychologia, 43(5), pp. 771–778.

82. Vallila-Rohter, S. and Kiran, S. (2013) ‘Non-linguistic learning and aphasia: Evidence from a paired associate and feedback-based task’, Neuropsychologia, 51(1), pp. 79–90.

83. Wechsler, D. (2008) Wechsler Adult Intelligence Scale (4th ed.). San Antonio, TX: Pearson Assessment.

84. Weiss, P. H., Ubben, S. D., Kaesberg, S., Kalbe, E., Kessler, J., Liebig, T., & Fink, G. R. (2016), ‘Where language meets meaningful action: A combined behavior and lesion analysis of aphasia and apraxia’, Brain Structure & Function, 221, pp. 563–576.

85. Wilke, M., de Haan, B., Juenger, H. and Karnath, H. O. (2011) ‘Manual, semi-automated, and automated delineation of chronic brain lesions: A comparison of methods’, Neuroimage, 56(4), pp. 2038–2046.

86. Yeung, O. and Law, S.-P. (2010) ‘Executive functions and aphasia treatment outcomes: Data from an ortho-phonological cueing therapy for anomia in Chinese’, International Journal of Speech-Language Pathology, 12(6), pp. 529–544.

87. Yourganov, G., Smith, K. G., Fridriksson, J. and Rorden, C. (2015) ‘Predicting aphasia type from brain damage measured with structural MRI’, Cortex, 73, pp. 203–215.

