## supplementary material for "Mapping language and non-language cognitive deficits in post-stroke anomic aphasia"

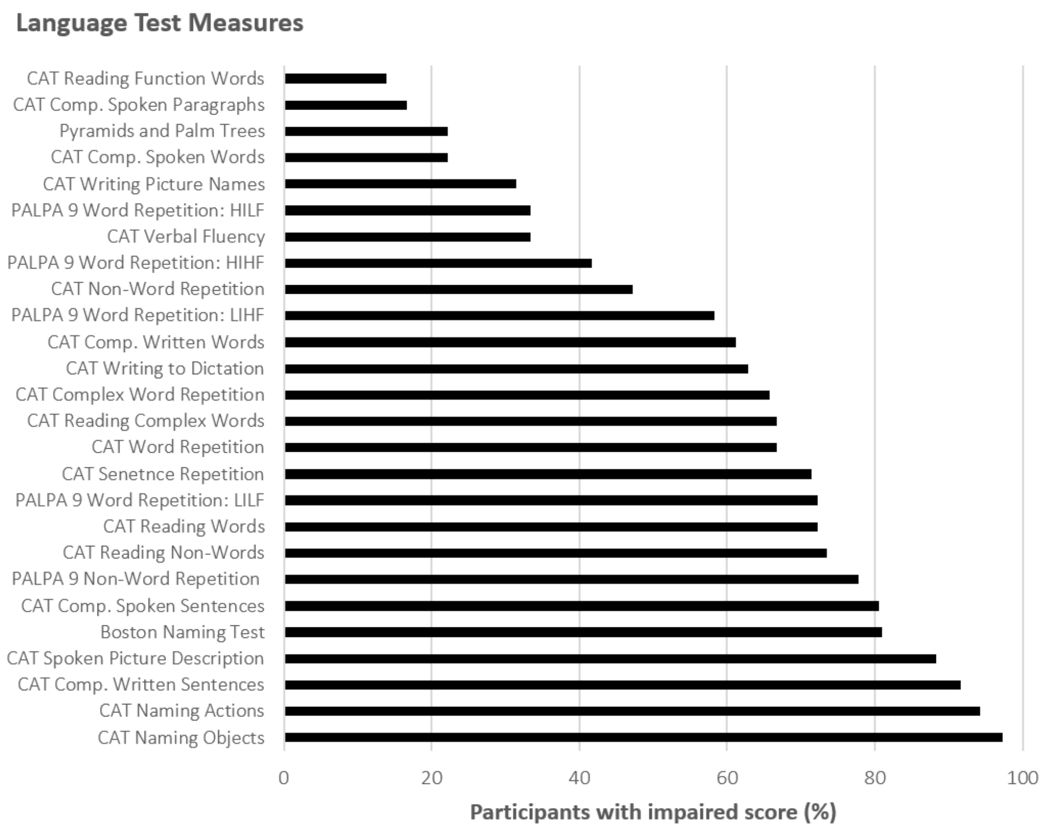


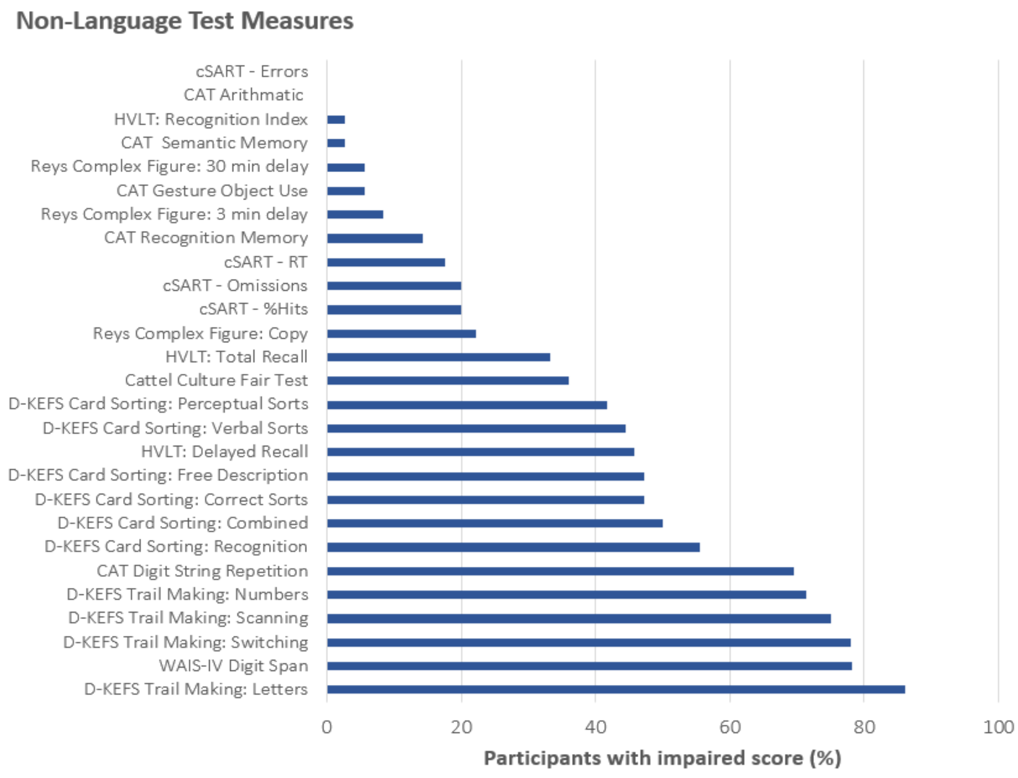


Supplementary Figure 1. Percentage of anomic participants with impaired performance on language and non-language tests. Impairment was determined based on a cut-off score defined by the test manual. Where this was not available, a cut-off was based on a score two standard deviations below the mean of a normative sample. Normative data for the Boston Naming test retrieved from Nicholas et al., 1989; Cattell's culture fare test (Tranter and Koutsaal, 2008). No cut-offs available for Raven's coloured progressive matrices and PALPA 8. See supplementary table 1 for individual participants' scores on behavioural assessments.

**Supplementary Table 1. Participants' scores on the behavioural assessments**

| | BNT | PALPA | | | | | | WAIS | | D-KEFS Card Sorting | | | | |
|---|---|---|---|---|---|---|---|---|---|---|---|---|---|---|
| **ID** | Boston Naming Test | PALPA 8 Repetition - Non-Words | PALPA 9 Repetition - Words (HIHF) | PALPA 9 Repetition - Words (HILF) | PALPA 9 Repetition - Words (LIHF) | PALPA 9 Repetition - Words (LILF) | PALPA 9 Repetition - Non-Words | Digit Span - Forwards | Digit Span - Backwards | Correct Sorts | Free Description | Recognition - Description | Verbal Sorts | Perceptual Sorts |
| 01 | 52 | 4 | **17** | **13** | **5** | **7** | **10** | 1 | 2 | 12 | 13 | 12 | 9 | 12 |
| 02 | **15** | 10 | **16** | **16** | **8** | **11** | **19** | 3 | 2 | 13 | 13 | 12 | 9 | 14 |
| 03 | 52 | 14 | **18** | **15** | 19 | **17** | **49** | 6 | 0 | **6** | **5** | **3** | 10 | **4** |
| 04 | **31** | 23 | 19 | **17** | **17** | **14** | **54** | 1 | 0 | 8 | 8 | 10 | **7** | 9 |
| 05 | 56 | 26 | 20 | 20 | 20 | 20 | 73 | 9 | 3 | 16 | 15 | 16 | 10 | 17 |
| 06 | **40** | 30 | 20 | 20 | 20 | 20 | 75 | 12 | 3 | 11 | 9 | 9 | **7** | 12 |
| 07 | **31** | 21 | 20 | 19 | **18** | 20 | 65 | 2 | 0 | 13 | 11 | **6** | 14 | 11 |
| 08 | **32** | 25 | **18** | 19 | 20 | **18** | **41** | 5 | 2 | **6** | **5** | **5** | **5** | **7** |
| 09 | **11** | 9 | **14** | **11** | **7** | **3** | **15** | 2 | 1 | **4** | **3** | **1** | **5** | **4** |
| 10 | **41** | 7 | **18** | **16** | **14** | **14** | **27** | 2 | 3 | **4** | **4** | **3** | **4** | **6** |
| 11 | **26** | 16 | **18** | 18 | **17** | **14** | **33** | 1 | 4 | **7** | **6** | **3** | 9 | **6** |
| 12 | **36** | 10 | **17** | **15** | **16** | **14** | **24** | 11 | 7 | 12 | 12 | 9 | 10 | 13 |
| 13 | 51 | 19 | 20 | 20 | 20 | **16** | **60** | 8 | 4 | 9 | 9 | 8 | 8 | 10 |
| 14 | 52 | 15 | **18** | 18 | **17** | **17** | **49** | 9 | 4 | 11 | 9 | 13 | 10 | 10 |
| 15 | 56 | 17 | 19 | 20 | **18** | **17** | **55** | 4 | 2 | 12 | 12 | 9 | 8 | 14 |
| 16 | 51 | 12 | 20 | 20 | 19 | 20 | **41** | 4 | 1 | 14 | 13 | 11 | 19 | 11 |
| 17 | **35** | 16 | 20 | **17** | **18** | **15** | **62** | 3 | 0 | 11 | 11 | 9 | 9 | 11 |
| 18 | **26** | 26 | 20 | 20 | 19 | **17** | 64 | 8 | 3 | **7** | **6** | **5** | 12 | **5** |
| 19 | **46** | 26 | 20 | 20 | 19 | 19 | 72 | 7 | 5 | 11 | 11 | 12 | **6** | 13 |
| 20 | **13** | 19 | 19 | 18 | **13** | **14** | **47** | 4 | 1 | **2** | **1** | **1** | **2** | **3** |
| 21 | **21** | 28 | 20 | 20 | 20 | 19 | 76 | 16 | 13 | 10 | 10 | 9 | 9 | 10 |
| 22 | **42** | 21 | 19 | 19 | **18** | 20 | **50** | 5 | 0 | **7** | **6** | **5** | 13 | **4** |
| 23 | **29** | 19 | **18** | **17** | **17** | **17** | **55** | 12 | 7 | 9 | 8 | 8 | 10 | 9 |
| 24 | **45** | 22 | **18** | 20 | **18** | 20 | **55** | 2 | 0 | 8 | 8 | **7** | **7** | 9 |
| 25 | **31** | 7 | **18** | 18 | **14** | **15** | **25** | 2 | 2 | 13 | 12 | **7** | 15 | 10 |
| 26 | **36** | 12 | 20 | 20 | 20 | **17** | **41** | 4 | 4 | **5** | **5** | **5** | **7** | **4** |
| 27 | **40** | 20 | **18** | 20 | **18** | **16** | **51** | 9 | 4 | **7** | **6** | **7** | 8 | **7** |
| 28 | **38** | 19 | 19 | 18 | **15** | **16** | **24** | 4 | 3 | 9 | 9 | 9 | **7** | 11 |
| 29 | **13** | 4 | 20 | 18 | **12** | **11** | **22** | 4 | 3 | **3** | **3** | **3** | **5** | **3** |
| 30 | **18** | 27 | 20 | **16** | 19 | **16** | 71 | 9 | 3 | **4** | **5** | **4** | **6** | 9 |
| 31 | **43** | 11 | **18** | **17** | **16** | **12** | **24** | 5 | 4 | **6** | **5** | **7** | 8 | **5** |
| 32 | **10** | 11 | **16** | **14** | 19 | **14** | **35** | 6 | 2 | **3** | **3** | **1** | **1** | **2** |
| 33 | **27** | 20 | 19 | 18 | **16** | **17** | **45** | 1 | 0 | **3** | **2** | **1** | **1** | **1** |
| 34 | **24** | 26 | 20 | 20 | 20 | **18** | **61** | 4 | 1 | **7** | **5** | **6** | **5** | 8 |
| 35 | **13** | 22 | 20 | 20 | 19 | 20 | **52** | 3 | 2 | **7** | **4** | **7** | **7** | **5** |
| 36 | **47** | 28 | 20 | 20 | 20 | 20 | 78 | 10 | 6 | 10 | 10 | 9 | 9 | 11 |
| **Max.** | 60 | 30 | 20 | 20 | 20 | 20 | 80 | 16 | 14 | 19 | 19 | 19 | 19 | 19 |
| **Cut-off** | 48 | NA | 18.61 | 17.66 | 18.61 | 18.51 | 62.5 | NA | NA | 7 | 7 | 7 | 7 | 7 |

### Supplementary Table 1 (cont.) Participants' scores on the behavioural assessments

| ID | CAT | | | | | | | | | | | | | | |
|---|---|---|---|---|---|---|---|---|---|---|---|---|---|---|---|
| | Comprehension- Spoken Words | Comprehension - Written Sentences | Repetition - Words | Repetition - Complex Words | Repetition - Non-Words | Repetition - Digit String | Repetition - Sentences | Naming - Objects | Naming - Actions | Spoken Picture Description | Reading - Words | Reading - Complex Words | Reading - Function Words | Reading - Non-Words | Writing to Dictation |
| 01 | **17** | **20** | **25** | **0** | **0** | **3** | **4** | **41** | **8** | **20** | 46 | **4** | 6 | **3** | **23** |
| 02 | 26 | **12** | **14** | **0** | **2** | **4** | **0** | **13** | **0** | **14** | **16** | **0** | 5 | **2** | **3** |
| 03 | **2** | **19** | **19** | **2** | **2** | 5 | 6 | **38** | **6** | **8** | 48 | 6 | 6 | **6** | **15** |
| 04 | 30 | **15** | **24** | **0** | 8 | **2** | **3** | **26** | **2** | **8** | **29** | **1** | 6 | **2** | **17** |
| 05 | 30 | **21** | **28** | 6 | 9 | 6 | 6 | **42** | **8** | 38 | 48 | 5 | 6 | 9 | 26 |
| 06 | 26 | **18** | 32 | 6 | 10 | 6 | 6 | **38** | **4** | 39.5 | 46 | 6 | 6 | **5** | 26 |
| 07 | 30 | **13** | 30 | 6 | 8 | **3** | **0** | **30** | **4** | **7** | **30** | **1** | 4 | **2** | **8** |
| 08 | **22** | **10** | **29** | **5** | **5** | **4** | **3** | **30** | **0** | **19** | **38** | 6 | 6 | 8 | **23** |
| 09 | **20** | **10** | **17** | **0** | **2** | **3** | **0** | **17** | **0** | **5** | **24** | **0** | **1** | **0** | **8** |
| 10 | 28 | 28 | **21** | **1** | **2** | **3** | **4** | **34** | **1** | **19.5** | **31** | **3** | 6 | **4** | 27 |
| 11 | 26 | **19** | **26** | 6 | 8 | **2** | **3** | **32** | **6** | **24** | **37** | **2** | 4 | 7 | **12** |
| 12 | **25** | **18** | **18** | **4** | **2** | 6 | **5** | **35** | **5** | **25.5** | **29** | **3** | 6 | **3** | **22** |
| 13 | 27 | **15** | 30 | 6 | 6 | 5 | **5** | **41** | **8** | **24** | 47 | **4** | 6 | **4** | 25 |
| 14 | 30 | **23** | **22** | **3** | 6 | 5 | 6 | **29** | **7** | **13** | **38** | **4** | 6 | 10 | 28 |
| 15 | 29 | **16** | 31 | **4** | 6 | **4** | **3** | **40** | **7** | **24** | 46 | 6 | 6 | 10 | **24** |
| 16 | 29 | **21** | 32 | 6 | 10 | **4** | **3** | **41** | **2** | **21.5** | **39** | **2** | 6 | 8 | **17** |
| 17 | 28 | **14** | **28** | **4** | 6 | **4** | **0** | **35** | **8** | **13** | **36** | **2** | 6 | **4** | **24** |
| 18 | 30 | **16** | **28** | **4** | 8 | 5 | 6 | **39** | **3** | **31.5** | 48 | **4** | 6 | 8 | 28 |
| 19 | 26 | 24 | 32 | 6 | 6 | **4** | **5** | **36** | **6** | **24.5** | **44** | 6 | 6 | **4** | **21** |
| 20 | **24** | **12** | **14** | **0** | **4** | **3** | **3** | **29** | **0** | **11** | **8** | **0** | 6 | **0** | **0** |
| 21 | 27 | **23** | 30 | 6 | 8 | 7 | 6 | **37** | **6** | **30.5** | 48 | 6 | 6 | 10 | 28 |
| 22 | 28 | **18** | 32 | 6 | 6 | **2** | **4** | **38** | **3** | **13** | **38** | **4** | 0 | **4** | **21** |
| 23 | 28 | **22** | **27** | **4** | 9 | 5 | 6 | **40** | 9 | **27** | 46 | 6 | 6 | **4** | 28 |
| 24 | 29 | **18** | **27** | **0** | 6 | **2** | **4** | **40** | **4** | **17.5** | **44** | **0** | 6 | **2** | **17** |
| 25 | 29 | **18** | **21** | **2** | **4** | **4** | **3** | **40** | **4** | **13** | **36** | **0** | 5 | **2** | 27 |
| 26 | 26 | **12** | 30 | 6 | **4** | **4** | **3** | **43** | **5** | 35 | **34** | **2** | 4 | **0** | **6** |
| 27 | 28 | **22** | 30 | **4** | **4** | 7 | 6 | **41** | **5** | 36.5 | 46 | 6 | 6 | 10 | 28 |
| 28 | 30 | **18** | **28** | **5** | 6 | **3** | 6 | **39** | 10 | **20** | **40** | 5 | 6 | **4** | 28 |
| 29 | 27 | **13** | **16** | **0** | **4** | **4** | **0** | **30** | **0** | **4** | **20** | **0** | **0** | **0** | **7** |
| 30 | **22** | **19** | **26** | **4** | 8 | **4** | 6 | **33** | **4** | **32** | **38** | 6 | 6 | **3** | **22** |
| 31 | 27 | **15** | **26** | **3** | 8 | **4** | **3** | **37** | **3** | **22** | **42** | **3** | 5 | **2** | 26 |
| 32 | **24** | **10** | **29** | 6 | **2** | **3** | **3** | **21** | **0** | **1** | **18** | **0** | **3** | **0** | **0** |
| 33 | 27 | **14** | **28** | **4** | **4** | **2** | **0** | **39** | **2** | **8** | **32** | **4** | 4 | **0** | **3** |
| 34 | **29** | **22** | **32** | **6** | **1** | **4** | **3** | **43** | **8** | NT | **39** | **4** | 6 | **2** | **12** |
| 35 | 28 | **15** | **26** | **0** | **4** | **2** | **3** | **29** | **3** | NT | **18** | **0** | **0** | **0** | **0** |
| 36 | 29 | 30 | 31 | 6 | 5 | 7 | 6 | 48 | **5** | **11** | **44** | 6 | 6 | 10 | 28 |
| **Max.** | 30 | 32 | 32 | 6 | 10 | 8 | 6 | 48 | 10 | _ | 48 | 6 | 6 | 10 | 28 |
| **Cut-off** | 25 | 23 | 29 | 5 | 5 | 4 | 5 | 43 | 8 | 33 | 45 | 4 | 3 | 6 | 24 |

This table displays individual participants' scores on the behavioural assessments that loaded on factors extracted from the rotated PCA. Scores below cut-off shown in bold. PALPA = Psycholinguistic Assessments of Language Processing in Aphasia; LILF = Low Intelligibility Low Frequency, LIHF = Low Intelligibility High Frequency, HIHF = High Intelligibility High Frequency, HILF = High Intelligibility Low Frequency. CAT = Comprehensive Aphasia Test. DKEFS = Delis-Kaplan Executive Function System. WAIS = Wechsler Adult Intelligence Scale. NA= data not available; NT= not tested.
